# *Trans*-cinnamaldehyde potently kills *Enterococcus faecalis* biofilm cells and prevents biofilm recovery

**DOI:** 10.1101/2020.08.09.243485

**Authors:** Islam A. A. Ali, Becky P. K. Cheung, Jukka. P. Matinlinna, Celine M. Lévesque, Prasanna Neelakantan

## Abstract

*Enterococcus faecalis* is a biofilm-forming, nosocomial pathogen that is frequently isolated from failed root canal treatments. Contemporary root canal disinfectants are ineffective in eliminating these biofilms and preventing reinfection. As a result, there is a pressing need to identify novel and safe antibiofilm molecules. The effect of short-term (5 and 15 min) and long-term (24 h) treatments of TC on the viability of *E. faecalis* biofilms was compared with currently used root canal disinfectants. Treatment for 15 min with TC reduced biofilm metabolic activity as effective as 1% sodium hypochlorite and 2% chlorhexidine. Treatment with TC for 24 h was significantly more effective than 2% chlorhexidine in reducing the viable cell counts of biofilms. This serendipitous effect of TC was sustained for 10 days under growth-favoring conditions. For the first time, our study highlights the strong antibacterial activity of TC against *E. faecalis* biofilms, and notably, its ability to prevent biofilm recovery after treatment.

## 1. Introduction

Surface-adherent communities of microorganisms, called biofilms, are implicated in a wide variety of human infections [1]. Biofilms consist of a complex, spatio-temporally organized extracellular matrix, which limits the penetration of antimicrobials, thereby rendering biofilm cells more tolerant to antimicrobials, compared to planktonic cells [2]. As such, the treatment of biofilms is considered a clinical challenge [3]. *Enterococcus faecalis* is a biofilm-forming pathogen that is implicated in a wide variety of pathological conditions including infective endocarditis, nosocomial, urinary tract and root canal infections [4].

Root canal treatment is a dental procedure performed to save infected teeth. Root canal asepsis is essential to establish an environment conducive to healing [5]. Furthermore, the conduit between the root canal and surrounding tissues easily causes spread of this infection [6]. Disinfection of the root canal attempted using antimicrobial agents applied during the clinical procedures to kill, and eradicate biofilms [7]. The commonly used current disinfectants for this short-term process are sodium hypochlorite and chlorhexidine [7]. In addition, interappointment dressings such as chlorhexidine are used to inactivate the residual microorganisms, and to maintain the aseptic environment until root canal filling [8]. There is evidence to show that these conventional disinfectants fail to achieve complete microbial killing and removal of root canal biofilms [9]. Notably, they are incapable of preventing biofilm recovery after treatment, which contributes to treatment failure [10, 11]. Although root canal infections are polymicrobial [12], *E. faecalis* is the most commonly isolated pathogen in failed treatments, and is thus considered the model organism to investigate the effect of novel therapies [13]. This is because of the remarkable activity of *E. faecalis* to survive and adhere to treated dentine surfaces [14], and its ability to tolerate nutrient-deprived environments encountered inside root canals [15]. That said, there is a critical need to develop alternative antimicrobials to improve root canal disinfection and prevent reinfection.

Plants are excellent sources of so-called essential oils (EOs), and secondary metabolites, which demonstrate a broad-spectrum antimicrobial activity and compatibility to host tissues [16]. Cinnamon is a native plant of China, India and South-East Asia [17]. *Trans*-cinnamaldehyde (TC), systematically called *trans*-3-phenyl-2-propenal (with the linear formula: C_6_H_5_CH=CHCHO), is a phenylpropanoid, which predominantly exists in cinnamon EOs [18]. It is Generally Recognized as Safe (GRAS) by the US Food and Drug Administration (US FDA) with well-known antibiofilm activity against pathogenic Gram-positive [19, 20] and Gram-negative bacteria [21, 22]. However, the effect of TC on *E. faecalis* biofilms remains to be investigated.

In this laboratory study, we investigated the effect of *trans*-cinnamaldehyde compared to conventional root canal disinfectants against *E. faecalis* biofilms. Furthermore, we tested and evaluated the ability of TC to prevent the recovery of *E. faecalis* biofilms under growth favouring conditions.

## 2. Materials and Methods

### 2.1. Bacterial strains and culture conditions

A clinical strain of *E. faecalis* isolated from failed root canal treatment (LAB846) was used in all the experiments. The strain cultured overnight in brain heart infusion (BHI) broth was washed twice in phosphate-buffered saline (PBS), and resuspended in BHI to 10^6^ CFU/ml. In all experiments, the biofilms were developed at 37°C for 72 h and the growth medium was replenished every 24 h. Our pilot study (data not shown) confirmed that these conditions resulted in the formation of a mature biofilm with abundant extracellular polymeric substance (EPS). All experiments were performed at three independent occasions in triplicates.

### 2.2. Chemicals

*Trans*-cinnamaldehyde (TC), dimethyl sulfoxide (DMSO), and chlorhexidine digluconate (CHX) were purchased from Sigma Aldrich (St. Louis, MO, USA). Six percent sodium hypochlorite (NaOCl) and 20% CHX solutions were diluted in sterile deionized water to obtain 1% NaOCl and 2% CHX working solutions. DMSO (5% v/v) was used as a solvent for TC [19]. Before use, all the chemicals were filter-sterilized by passing through 0.2 μm-pore syringe filters (Pall Life Sciences, Pensacola, FL, USA).

### 2.3. Short-term treatment of E. faecalis biofilms

The effects of TC on biofilm viability were compared and contrasted to 1% NaOCl and 2% CHX. Briefly, biofilms were developed in sterile 96-well polystyrene plates for 72 h. After incubation, the planktonic supernatant was carefully aspirated, and the biofilms were washed with PBS to remove the non-adherent cells. The biofilms were then treated with TC (0.5, 0.75 and 1%), NaOCl, CHX, DMSO or deionized water for 5 and 15 min. The treatment solutions were then carefully aspirated, biofilms were rinsed with PBS and the biofilm viability was evaluated using the XTT metabolic activity as described previously [23]. Briefly explained, 200 μl of (XTT + menadione) reaction solution was added to each well, and incubated in dark for 3 h at 37°C. After incubation, 100 μl of the supernatant was transferred to a new microplate and the optical density was measured at 492 nm (SpectraMax M2; Molecular Devices, Sunnyvale, CA, USA).

### 2.4. Long-term treatment of E. faecalis biofilms

#### 2.4.1. Biofilm viability: Immediate and sustained post-treatment effects

Seventy-two hours old biofilms were developed in sterile 96-well polystyrene plates as described above. The biofilms were treated with TC (0.5 and 1%), 2% CHX, DMSO or deionized water for 24 h. The effect on biofilm viability was evaluated immediately after treatment (D_0_). In parallel, another set of treated biofilms were further incubated in BHI with daily replenishment of growth media. On the 10^th^ day after treatment (D_10_), biofilm viability was evaluated. At the two time points (D_0_ and D_10_), the biofilm viability was evaluated using the XTT-metabolic activity (as described above), and CFUs assays. Next, the biofilms were collected by vigorous scraping and pipetting. The biofilm suspensions were serially-diluted, plated on blood agar, and the colonies were counted after of incubation at 37°C for 48 h.

#### 2.4.2. Microscopic imaging of biofilms: Biofilm viability and morphology

Biofilms were developed on sterile collagen-coated hydroxyapatite (HA) discs (Clarkson Chromatography Products, Williamsport, PA, USA) in sterile 24-well polystyrene plates based on a previously described protocol [24]. At the end of the incubation period, the discs were washed in PBS to remove the non-adherent cells, transferred to new wells containing 0.5% TC, 2% CHX, or deioinized water, and further incubated for 24 h. The biofilms were stained using LIVE/DEAD Baclight viability kit (L-7012, Molecular Probes, Eugene, OR, USA) containing SYTO-9 and propidium iodide (PI) for 15 min following the manufacturer’s instructions. The biofilms were observed using an oil-immersion objective lens (x 100) of a confocal laser scanning microscope (CLSM; FLUOVIEW FV1000, Olympus, Tokyo, Japan) at five randomly selected points. The biofilm z-stacks were analyzed to quantify the percentage of the apparently dead biofilm cells. Reconstructed 3D images were analyzed to evaluate the biofilm architecture (Imaris, Bitplane, St. Paul, MN, USA).

The effect of TC and CHX on the morphology of *E. faecalis* biofilms was assessed using scanning electron microscopy (SEM). The biofilms were processed as described previously [24]. The biofilm morphology was evaluated at randomly selected points on each HA disc.

#### 2.4.3. Water-insoluble exopolysaccharides

The water-insoluble exopolysaccharides of the treated biofilms were quantified using the phenol-sulfuric acid method [25]. The biofilms were washed with PBS and collected by scraping and vigorous pipetting. The suspensions were homogenized by vortexing and sonication and centrifuged for 5 min at 10,000 rpm. The pellets were resuspended in sterile deionized water. Aliquots of the biofilm suspension were mixed with 5% phenol and 98% concentrated sulfuric acid, and incubated at 90°C for 30 min. The optical density was measured at 492 nm. The concentration of exopolysaccharides was determined using a standard curve generated using glucose as a standard.

### 3. Statistical analysis

All data are presented as mean ± standard deviation. One-way or two-way ANOVA with the *post hoc* Bonferroni test was used to compare the results between the experimental groups. The data were analyzed using SPSS (Version 25, Chicago, IL, USA). *P* < 0.05 was considered as statistically significant.

## 4. Results

### 4.1. Short-term treatment of E. faecalis biofilms

The effects of *trans-*cinnamaldehyde, NaOCl and CHX on the metabolic activity of 72 h-old *E. faecalis* biofilms were evaluated using the XTT assay. The treatment durations were based on clinically relevant time frames during root canal treatment. After treatment for 5 min, TC significantly reduced the metabolic activity of biofilms compared to the control (*P* < 0.001, Figure 1), but was significantly less effective compared to NaOCl and CHX (*P* ≤ 0.001, Figure 1). When the biofilms were treated for 15 min, the metabolic activity was significantly reduced in all treatment groups (*P* < 0.001, Figure 1), and no significant difference was observed between TC, NaOCl, and CHX treatment (*P* > 0.05, Figure 1).

**Figure 1.**
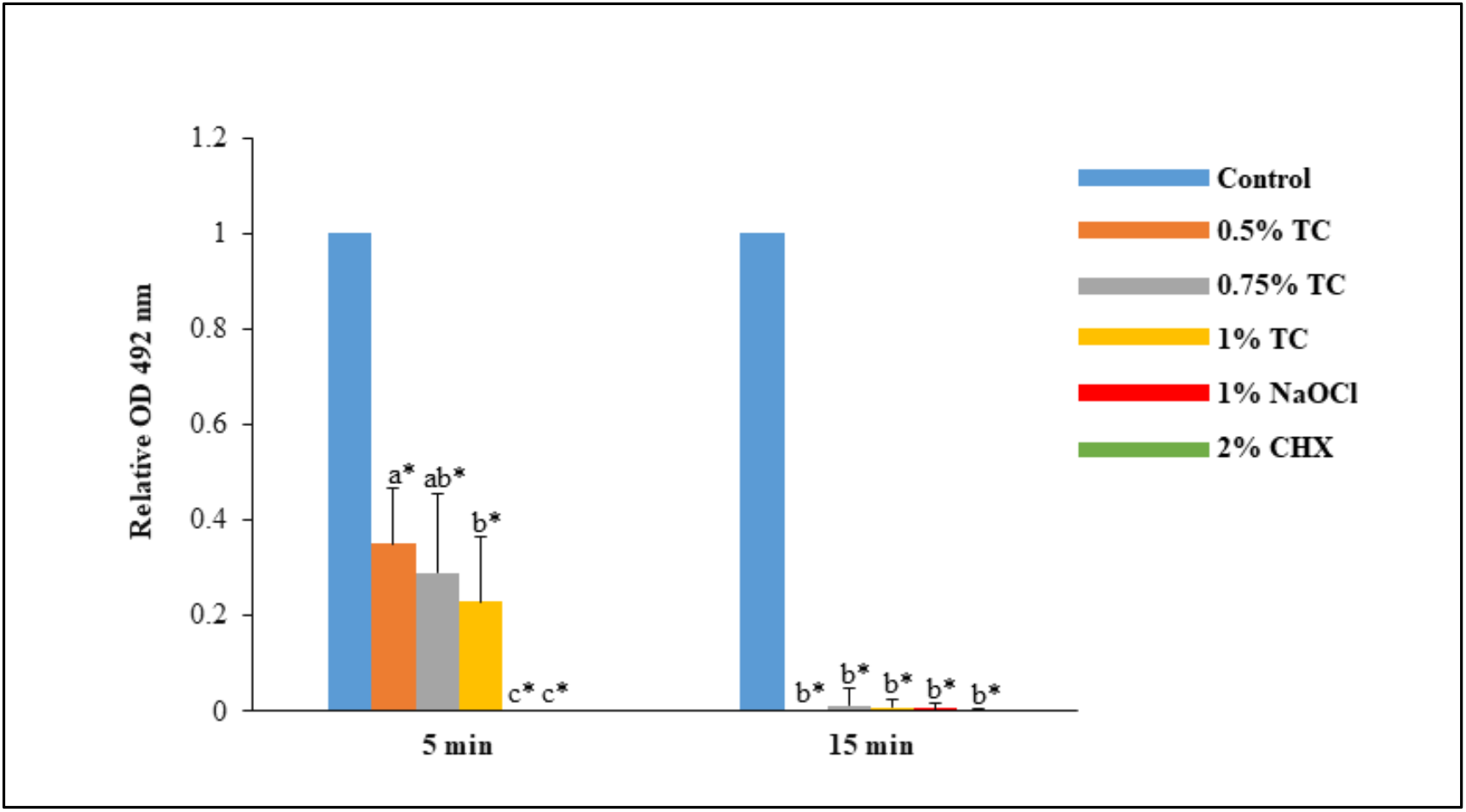
Metabolic activity of 72 h-old *E. faecalis* biofilms after treatment for 5 and 15 min. OD_492_ readings in TC, NaOCl and CHX groups were expressed relative to the control. Different alphabets indicate a statistically significant difference between treatment groups. * indicates significant difference between each treatment group and the control (*P* < 0.05 was considered as statistically significant).

### 4.2. Long-term treatment of E. faecalis biofilms

Metabolic activity and CFUs assays were employed to assess the immediate and sustained post-treatment effects of TC (0.5, 1%), CHX, DMSO or deionized water on *E. faecalis* biofilm viability. Immediately after treatment (D_0_), TC and CHX significantly diminished the metabolic activity compared (*P* < 0.001, Figure 2A). TC treatment resulted in significantly lower log_10_ CFU/ml compared to CHX and water (*P* < 0.01, Figure 2B). After incubation of treated biofilms in BHI for 10 days, biofilms treated with TC or CHX did not show any significant increase in the metabolic activity at D_10_ compared to D_0_ (*P* > 0.05, Figure 2A). However, there was no significant change in the log_10_ CFU/ml of TC-treated biofilms at D_10_ compared to D_0_ (*P* > 0.05, Figure 2.), whereas in CHX group, the log_10_ CFU/ml at D_10_ was significantly greater than at D_0_ (*P* < 0.001, Figure 2B).

**Figure 2.**
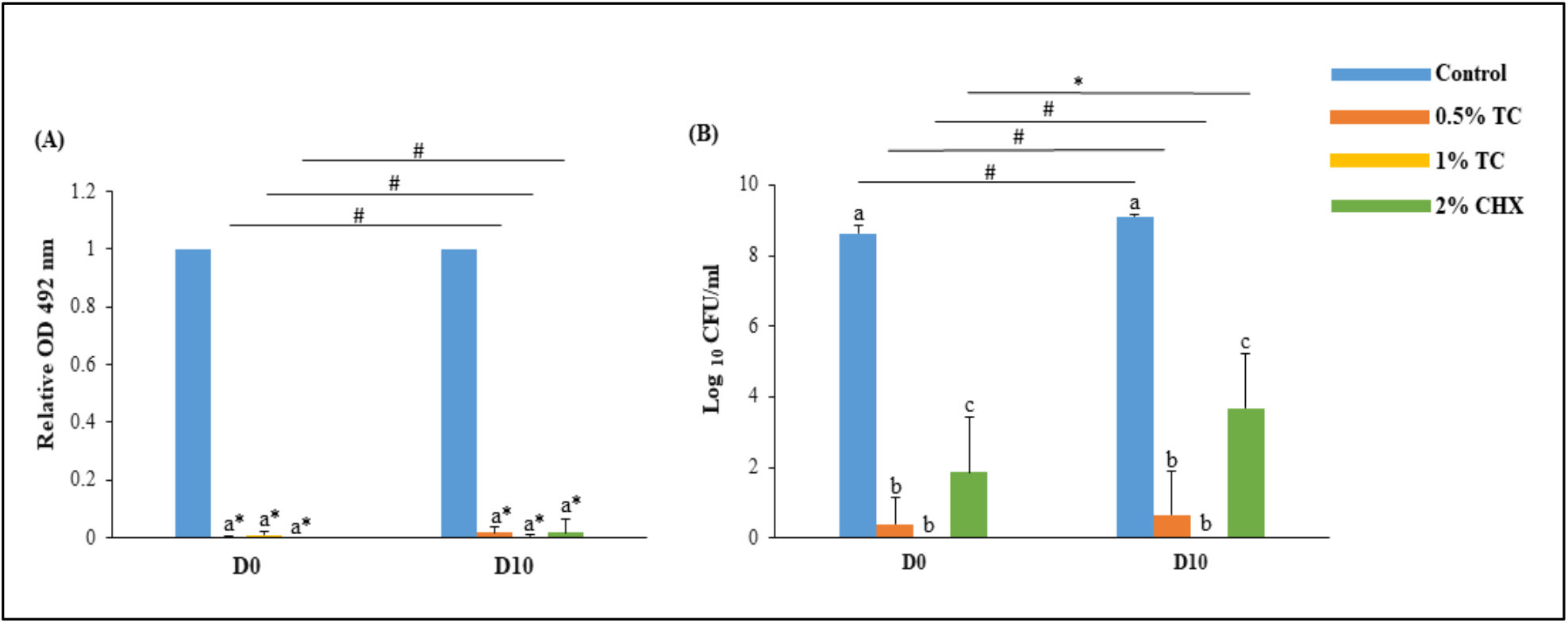
The viability of 72 h-old *E. faecalis* biofilms immediately after treatment for 24 h (D_10_), and after incubation in brain heart infusion (BHI) broth (D_0_). The viability was investigated using the (A) XTT metabolic activity assay: The OD 492 nm readings in TC and CHX groups were expressed relative to the control. * indicates significant difference between each treatment group and the control, while # indicates no significant difference between D_0_ and D_10_. (B) Colony Forming Units: Changes in log_10_ CFU/ml between the D_0_ and D_10_ is shown for each experimental group. * and # indicate significant and non-significant differences respectively (*P* < 0.05 was considered as statistically significant).

We then assessed the viability of *E. faecalis* biofilm cells using fluorescent live/dead viability assay. Quantitative analysis revealed a significantly higher percentage of dead cells in *trans-*cinnamaldehyde-treated biofilms (95 ± 3.1%) compared to CHX (33 ± 11.2 %), and deionized water (2 ± 1.5 %) (*P* < 0.001, Figure 3A). The post-treatment changes in biofilm morphology were also observed using SEM. The control biofilms were composed of dense, closely aggregated cell clusters covering the entire surface of the HA discs. In biofilms treated with TC or CHX, the overall structure and morphology appear largely unaltered (Figure 3B).

**Figure 3.**
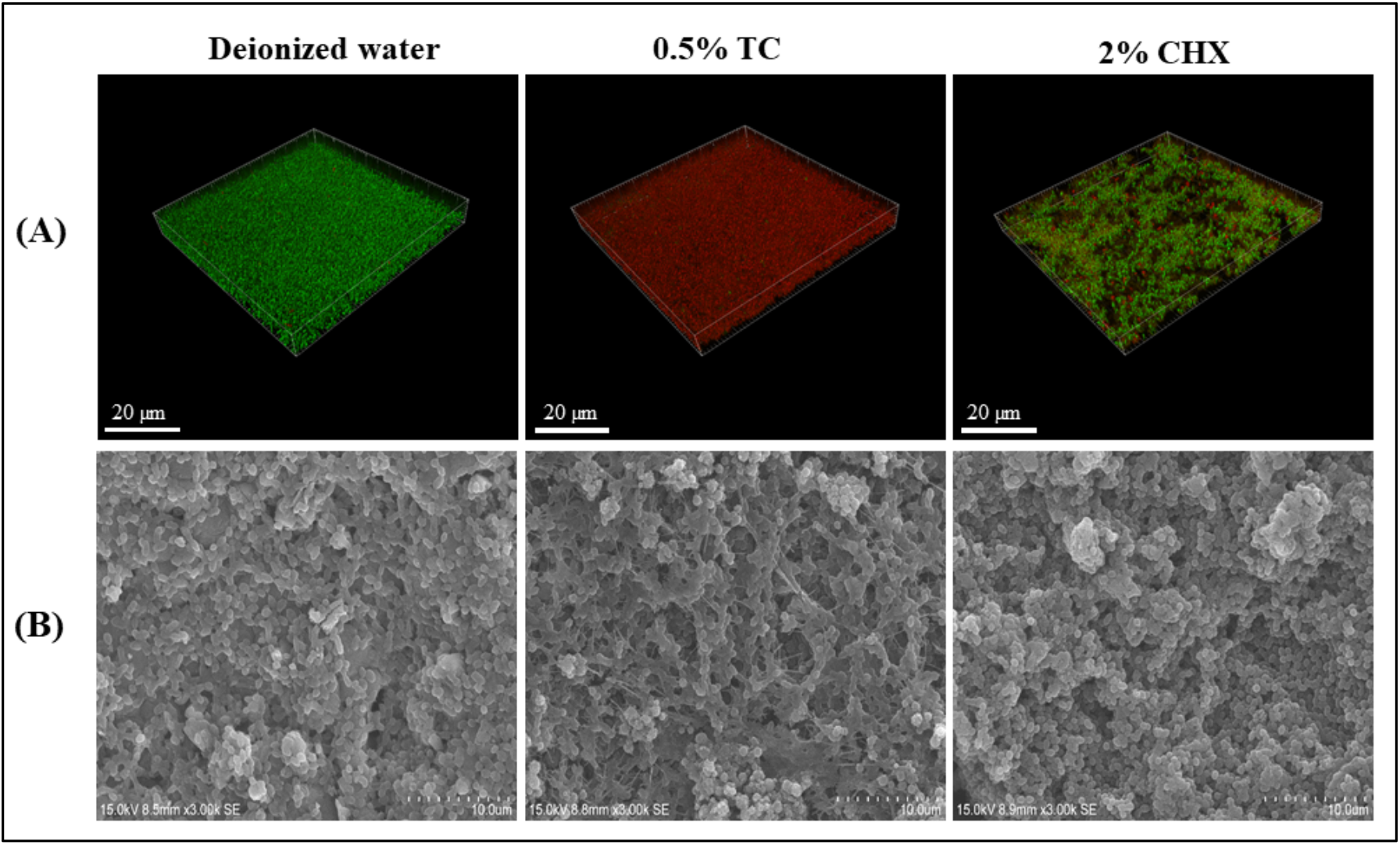
(A) Confocal microscopic images (x 100) and SEM images (x 3000) of 72 h-old *E. faecalis* biofilms on collagen-coated HA after treatment with deionized water, 0.5% TC and 2% CHX for 24 h. In CLSM images, the green fluorescent dye (STYO-9) labels cells with intact cell membranes, while the red fluorescent dye (PI) labels cells with damaged membranes. Deionized water and TC-treated biofilms are primarily composed of apparently live and dead cells respectively, while CHX-treated biofilms are composed of apparently dead (labelled red), partially damaged (labelled yellow or yellowish green) and live (labelled green) cells. The yellow or yellowish green-labelled cells are due to partially damaged cell membranes; thus SYTO9/PI are retained within the cells [26]. (B) SEM images reveals unaltered biofilm morphology after treatment with TC or CHX compared to deionized water.

The effect of TC on water-insoluble exopolysaccharides was evaluated using the phenol-sulfuric acid method. After treatment for 24 h, the exopolysaccharide content in TC-treated biofilms (223 ± 48 μg/ml) was not significantly different compared to deionized water (192 ± 75 μg/ml; *P* > 0.05). By contrast, treatment with CHX significantly decreased the exopolysaccharides content (94 ± 54 μg/ml) compared to TC- and deionized water-treated biofilms (*P* < 0.001).

## 5. Discussion

Natural plants are rich sources of essential oils and secondary metabolites, which protect them against pathogens and competitors in their natural environments [27]. In the current study, we investigated the effect of *trans-*cinnamaldehyde, the main constituent of cinnamon oil, on *E. faecalis* biofilms to explore its incorporation in therapeutic strategies. We used a clinical strain of *E. faecalis* isolated from failed root canal treatment. It has been importantly demonstrated that biofilms of clinical strains show an increase in the expression of surface proteins, rendering them less amenable to killing and removal. This has been attributed to its exposure *in vivo* to root canal disinfectants, and survival in nutrient-deprived environment [28]. Given that, our results have profound translational significance.

The investigations were performed after short-, and long-term exposure to TC to simulate its use as a root canal irrigant, and an interappointment dressing respectively. The concentrations of TC used in the current study were based on our preliminary experiments, which demonstrated killing of *E. faecalis* planktonic cells at 0.4% (data not shown). Our results demonstrated that *trans-*cinnamaldehyde reduced the viability of mature *E. faecalis* biofilms compared to deionized water, with just 5 min of exposure, but was less effective than NaOCl and CHX. When the treatment was extended to 15 min, TC diminished the biofilm viability as effective as NaOCl and CHX. The killing effect of TC is attributed to its ability to penetrate and perturb the integrity of hydrophobic cell membrane of *E. faecalis*. Compromised membrane integrity results in leakage of intracellular contents, and inhibition of the membrane–bound ATPase activity [29]. The *trans-*cinnamaldehyde molecule possesses the acrolein group (α, β-unsaturated carbonyl group) which is thought to be essential for its antibacterial activity [30].

Biofilm recovery after antimicrobial treatment is a critical cause of persistent and chronic infections. Following antimicrobial challenge, residual biofilm microbes are able to survive under harsh conditions [31]. Upon exposure to favourable growth conditions and depletion of residual antimicrobials, these cells revert into a metabolically active status, form new biofilms and induce post-treatment disease [32]. We have questioned whether *E. faecalis* biofilm cells treated with TC (0.5, 1%) and 2% CHX could recover under growth favouring conditions. Therefore, we investigated the immediate and sustained post-treatment effects of TC and CHX on *E. faecalis* biofilms. Notably, we found an increase in CFUs in CHX-treated biofilms at D_10_ compared to D_0_, despite the previously reported substantivity of CHX [11]. By contrast, and for the first time, our results indicate that TC prevented the recovery of *E. faecalis* biofilms as there was no significant increase in CFUs at D_10_ compared to D_0_. Despite the significant increase in viable colonies recovered from CHX-treated biofilms compared to TC at D_10_, the XTT assay showed no significant difference. This might be attributed to the minimum cell density (4.5-5 log_10_ CFU/ml) of metabolically active cells that is required to allow enzymatic reduction in tetrazolium salts-based assays [33]. In TC and CHX groups, the viable cell count at D_10_ remained below this threshold, indicating an important methodological issue with the XTT assay.

The presence of viable cells immediately after treatment in CHX-treated biofilms is in accordance with Li et al. [34]. Complete killing of *E. faecalis* biofilm cells by CHX was previously reported [35]. Differences in the strains, treatment time and experimental model could explain the different results. Here, we used a clinical strain, which formed prolific biofilms *in vitro* (data are not shown). Recovery in CHX-treated biofilms has been reported in previous studies [11, 36], in which the proportion of live cells declined for a week after treatment, and then gradually increased beyond the first week. The electrostatic attraction of CHX to EPS likely limits its penetration, and reduces its concentration to sublethal levels in deep biofilm layers [11]. The profound effect of TC in preventing biofilm recovery compared to CHX, might be due to a significantly higher population of dead cells after TC treatment. In addition, TC attacks crucial physiological processes, that could assist in biofilm recovery such as cell division, ATP generation system, diffusion of ions and metabolites, and quorum sensing [29]. Mechanistic studies on recovered biofilm cells may explain the results reported here, and that is beyond the scope of this study.

The structural changes in the treated biofilms were visualized on collagen-coated HA discs under CLSM and SEM. The compromised cell membrane integrity of TC-treated *E. faecalis* reflected in our CLSM findings, as the majority of biofilms cells were stained red, unlike the intact green-labelled cell population in the control biofilms. In biofilms treated with 2% CHX, a mixed population of green, yellowish green and red-labelled cells was observed indicating that CHX is less effective against biofilm cells. Our CLSM findings are in agreement with previous studies [37, 38], which showed apparently live cells after 24 h treatment with 2% CHX. The limited ability of CHX to disrupt attached biofilms is well known [39], and could be attributed to its potential fixative action [40]. This may also be explained by the upward relocation of EPS [41], which render the cells in deep biofilm layers more tolerant to CHX. TC and CHX-treated biofilms showed an unaltered morphology as demonstrated by SEM. Both treatments target the cell membrane integrity [18, 42]. Cytoplasmic coagulation, denaturation and rigidity of bacterial cell membranes in response to TC, render alterations in biofilm morphology less likely [43].

On the other hand*, trans-*cinnamaldehyde was unable to reduce the exopolysaccharides content significantly compared to the control, revealing that TC primarily targets the biofilm cells without degrading the matrix. Reduction of exopolysaccharides in CHX-treated biofilms is likely due to the contraction of polysaccharides chains of EPS caused by the electrostatic attraction of the cationic CHX molecules to the negatively charged EPS matrix [11]. Although the biofilm matrix contains polysaccharides, proteins, nucleic acids and lipids [2], we quantified only the polysaccharides as they contribute to the biofilm attachment to surfaces, mechanical stability, resistance to host defences, tolerance to antimicrobials and nutritional supply to biofilm bacteria [2].

In the results of the current study, we provide the first evidence on the antibiofilm effects of *trans-*cinnamaldehyde on *E. faecalis*, compared to traditional antimicrobials used in root canal treatment. Despite the fact that root canal biofilms are polymicrobial, the current study shed significantly new light on TC as a potential antimicrobial agent against *E. faecalis*, a major pathogen of persistent root canal infections. Future studies will be performed on multispecies biofilms in clinically relevant models and under different environmental conditions. Furthermore, the results of this study will have profound implications in chronic infections caused by *E. faecalis* elsewhere in the body.

## 6. Conclusions

- *Trans*-cinnamaldehyde kills *E. faecalis* biofilm cells as effectively as 1% sodium hypochlorite and 2% chlorhexidine after treatment for 15 min.
- *Trans*-cinnamaldehyde sustained its effect on *E. faecalis* biofilms under growth favouring conditions, in contrast to chlorhexidine, which demonstrated viable cells 10 days after treatment.

## Acknowledgments

This study was supported by the seed fund from the University of Hong Kong to Prasanna Neelakantan. Islam A. A. Ali is supported by the postgraduate fellowship from the University of Hong Kong, and this study is a part of his doctoral thesis.

## CRediT authorship contribution statement

**Islam A. A. Ali:** Conceptualization, methodology, formal analysis, investigation, writing, and visualization. **Becky P. K. Cheung:** Investigation and writing. **Jukka. P. Matinlinna:** Writing, and supervision, **Celine M. Lévesque:** Conceptualization, methodology, writing, and supervision**. Prasanna Neelakantan:** Conceptualization, methodology, writing, visualization, supervision, project administration, and funding acquisition.

